# Gain of Function Analysis Reveals Non-Redundant Roles for the *Yersinia pestis* Type III Secretion System Effectors YopJ, YopT, and YpkA

**DOI:** 10.1101/310243

**Authors:** Samantha G. Palace, Megan K. Proulx, Rose L. Szabady, Jon D. Goguen

**Author notes:** Address correspondence to Jon D. Goguen. Present addresses:Samantha G. Palace: Department of Immunology and Infectious Diseases, Harvard T. H. Chan School of Public Health, Boston, Massachusetts, USA Rose L. Szabady: Vedanta Biosciences, Inc., Cambridge, Massachusetts, USA.

## Abstract

Virulence of *Yersinia pestis* in mammals requires the type III secretion system, which delivers seven effector proteins into the cytoplasm of host cells to undermine immune responses. All seven of these effectors are conserved across *Y. pestis* strains, but three – YopJ, YopT, and YpkA – are apparently dispensable for virulence. Some degree of functional redundancy between effector proteins would explain both observations. Here, we use a combinatorial genetic approach to define the minimal subset of effectors required for full virulence in mice following subcutaneous infection. We found that a *Y. pestis* strain lacking YopJ, YopT, and YpkA is attenuated for virulence in mice, and that addition of any one of these effectors to this strain increases lethality significantly. YopJ, YopT, and YpkA likely contribute to virulence via distinct mechanisms. YopJ is uniquely able to cause macrophage cell death *in vitro* and to suppress accumulation of inflammatory cells to foci of bacterial growth in deep tissue, whereas YopT and YpkA cannot. The synthetic phenotypes that emerge when YopJ, YopT, and YpkA are removed in combination provide evidence that each enhances *Y. pestis* virulence, and that YopT and YpkA act through a mechanism distinct from that of YopJ.

## Introduction

*Yersinia pestis*, causative agent of plague, is notorious for its role in the European Black Death pandemics of the Middle Ages. Its pathogenesis in the mammalian host is remarkable: following inoculation of a small number of bacteria in the dermis by the bite of an infected flea, *Y. pestis* rapidly invades distal tissues and the vasculature. The resulting dense bacteremia enhances transmission, as it allows colonization of naïve fleas that ingest a sub-microliter blood meal. Dissemination of *Y. pestis* from the dermis to the bloodstream requires several bacterial adaptations that work in concert to achieve near-absolute suppression of the innate immune responses that would otherwise contain or clear the infection.

The *Y. pestis* type III secretion system (T3SS) is a major contributor to this innate immune evasion strategy. The T3SS transports bacterial proteins, called effectors, into the cytoplasm of target eukaryotic cells (1, 2). The bacterial translocon proteins YopB and YopD form a pore in the host cell membrane and interact with the syringe-like T3SS “injectisome” apparatus. This assembly is thought to form a continuous conduit that transports effector proteins directly from the intracellular compartment of the bacterial cell into the cytosol of target cells (2–4), though some recent data has challenged this model (5, 6).

The T3SS of the pathogenic yersiniae targets innate immune cells *in vivo* (7) and undermines a variety of antimicrobial responses in these cells, including phagocytosis, immune signaling, and the production of reactive oxygen species (ROS) (reviewed in (8)). Intoxication of these cells by the T3SS is one of the most important mechanisms underlying the innate immune evasion that is so crucial for *Y. pestis* virulence, and spontaneous loss of the pCD1 plasmid that encodes the T3SS profoundly attenuates *Y. pestis* in mammalian infection models (9–11). Mutations that compromise the type III secretion mechanism by inactivating injectisome components are likewise highly attenuating (12, 13).

*Y. pestis* shares a conserved set of seven T3SS effectors with the enteropathogenic yersiniae *Y. enterocolitica* and *Y. pseudotuberculosis*. Four of these effectors – YopH, YopE, YopK, and YopM – are required for full virulence of *Y. pestis* in murine infection models (14–20), although the attenuation associated with YopM deletion seems to vary among strains (21). YopH, YopE, and YopM directly target innate immune responses: YopH and YopE inhibit the production of reactive oxygen species (ROS) (22, 23) and interfere with phagocytosis (24–27), while YopM likely enhances virulence by preventing caspase 1 signaling (28, 29) and pyrin inflammasome activation (30, 31). The attenuation of YopK mutants may result from dysregulated secretion of the other effector proteins and the translocon proteins (19, 32).

The effector YopJ profoundly deranges host cell death signaling pathways *in vitro*, and as a result YopJ has been intensively studied in all three pathogenic *Yersinia* species. YopJ induces caspase-8/RIP-1 mediated apoptosis in macrophages, inhibits transcription of pro-inflammatory cytokines by **NFκB**, and may also stimulate caspase-1 signaling (33–35). However, *Y. pestis* mutants lacking the *yopJ* gene have been shown more than once to retain full virulence *in vivo* (36, 37). Single knock-outs of the cysteine protease YopT and the serine/threonine kinase YpkA have not been reported to impact virulence of *Y. pestis* in mammalian infection models.

Although YopJ, YopT, and YpkA are individually dispensable for *Y. pestis* virulence, all three effectors are conserved across natural and experimental *Y. pestis* strains. Given the small number of T3SS effectors found in *Y. pestis*, especially compared with the T3SSs of many other Gram-negative pathogens, it is likely that these effectors are selectively maintained because they play a role during some stage of the natural *Y. pestis* transmission cycle. However, these three effectors share some putative targets with one another and with the other T3SS effectors at both the protein and pathway level (8). Their functions may therefore overlap sufficiently to account for the observation that single deletion of any one does not have a measurable effect, at least in the context of standard laboratory survival studies using inbred mouse strains.

Traditional single gene knockout models are ill-suited to studying the individual contributions of the T3SS effectors, which share a high degree of interconnectedness among their putative target molecules and pathways. For example, no fewer than four effectors (YopH, YopE, YpkA, and YopT) are reported to interfere with phagocytic function of innate immune cells (reviewed in (8)). For this reason, if a mutation in a single effector fails to yield an attenuation phenotype *in vivo*, it is difficult to distinguish between true dispensability of the effector’s function and possible functional redundancy with other components of the T3SS.

To determine the effects of the YopJ, YopT, and YpkA effectors, we chose instead to test for gain-of-function phenotypes when effectors were added back to a strain from which all seven effector proteins had been deleted. This combinatorial genetic approach focuses on finding synthetic phenotypes, and is therefore robust to functional redundancy. We were able to dissect the individual contribution of each effector to pathogenesis in the intact animal. All seven effector proteins enhance virulence in mice – the first clear demonstration that YopJ, YopT, and YpkA can each contribute to *Y. pestis* virulence *in vivo*. The collection of combinatorial effector knock-outs we generated also allowed us begin the process of quantifying and characterizing the non-redundant role that each effector plays during infection. While it seems likely that the contribution of YopJ results from its effective killing of macrophages, as has been reported, the mechanisms underlying the contributions of YopT and YpkA remain uncertain despite considerable knowledge of their biochemical activity.

## Results

### Effectors YopH, YopE, YopK, and YopM are not sufficient for full virulence *in vivo*

To understand the functional role of each of the *Y. pestis* T3SS effector proteins during infection, we set out to find the minimal subset of effectors that were sufficient to mediate full virulence through the subcutaneous route of infection.

We approached this problem by first constructing KIM1001ΔT3SE, an unmarked strain that retains the T3SS injectisome, regulatory elements, and translocon proteins, but carries in-frame deletions in the open reading frames (ORFs) for all seven effector proteins (see Table S1 for details). This strain, constructed in the fully-virulent background KIM1001 (38), was highly attenuated through the subcutaneous route of infection. When infected with 10^3^ CFU KIM1001ΔT3SE, 0 of 8 mice developed visible symptoms of disease, and all mice survived infection (Table 1).

**Table 1.** Virulence of *Y. pestis* strains expressing subsets of type III secretion effector proteins. All infection experiments were performed with a subcutaneous dose of 1x10^3^ CFU in C57BL/6 mice. Median time to death references only those mice within each group that succumbed to infection; any mice that survived the duration of the experiment were excluded from the calculation of this metric. Significance was calculated for each curve compared to the KIM1001 curve by the Mantel-Cox test.

| Strain | Percent lethality<br>(number of mice<br>killed/total number<br>of mice infected) | Median<br>time to<br>death<br>(days) | Significance<br>against<br>KIM1001<br>(p value) |
| --- | --- | --- | --- |
| KIM1001ΔT3SE | 0% (0/8) | NA | < 0.0001 |
| KIM1001ΔT3SE::+yopHE | 0% (0/10) | NA | < 0.0001 |
| KIM1001ΔT3SE::+yopHEM | 0% (0/6) | NA | < 0.0001 |
| KIM1001ΔT3SE::+yopHEK | 0% (0/6) | NA | < 0.0001 |
| KIM1001ΔT3SE::+yopHEKM | 53% (9/17) | 9 | < 0.0001 |
| KIM1001ΔT3SE::+yopHEKMA | 88% (14/16) | 6 | 0.1305 |
| KIM1001ΔT3SE::+yopHEKMT | 87% (13/15) | 6 | 0.1184 |
| KIM1001ΔT3SE::+yopHEKMJ | 85% (22/26) | 5.5 | 0.1031 |
| KIM1001ΔT3SE::+yopHEKMJT | 100% (5/5) | 7 | 0.1501 |
| KIM1001ΔT3SE::+yopHEKMAJ | 100% (5/5) | 7 | 0.4756 |
| KIM1001ΔT3SE::+yopHEKMAT | 80% (4/5) | 7 | 0.1206 |
| KIM1001ΔT3SE::+yopHEKMAJT | 100% (15/15) | 5 | 0.3890 |
| KIM1001 | 100% (12/12) | 4.5 | NA |

Functional ORFs for various effectors were restored at their original loci in the ΔT3SE genetic background to generate strains expressing defined subsets of effectors. YopH and YopE are more strictly required for bacterial fitness *in vivo* than any of the other effector proteins (18). However, we found that these two effectors in combination are not sufficient for virulence. The strain KIM1001ΔT3SE::+yopHE, expressing YopH and YopE but carrying in-frame deletions in all of the remaining five effectors’ ORFs, failed to sicken or kill any mice following subcutaneous infection with 10^3^ CFU (n=10). By contrast, this dose is uniformly fatal with wild-type KIM1001 (Table 1).

Single deletion studies have demonstrated that YopM and YopK are also essential for full virulence of *Y. pestis* (19, 20). However, neither of these effectors was sufficient to restore virulence in the KIM1001ΔT3SE::+yopHE background. Strains expressing only YopH, YopE, and YopM (KIM1001ΔT3SE::+yopHE**M**) or YopH, YopE, and YopK (KIM1001ΔT3SE::+yopHE**K**) remained attenuated (0 out of 6 mice killed for each group following subcutaneous infection with 10^3^ CFU) (Table 1), though 2 out of 6 mice infected with KIM1001ΔT3SE::+yopHE**K** lost their fur around the injection site and developed local redness of the skin that persisted for at least 28 days.

The KIM1001ΔT3SE::+yopHE**KM** strain expresses all effectors previously reported to be necessary for full virulence, as assessed by subcutaneous infection based of single knockout mutants. This strain was substantially more virulent than the previous strains expressing subsets of these effectors, but significantly attenuated relative to the wild-type strain. KIM1001ΔT3SE::+yopHEKM killed approximately 50% of infected mice (9 out of 17) at a dose of 10^3^ CFU, indicating a ~100-fold increase of LD_50_ compared to wild-type KIM1001. Simultaneously restoring functional copies of the *ypkA, yopJ*, and *yopT* ORFs to this strain (generating the strain KIM1001ΔT3SE::+yopHEKMAJT, genetically identical to the wild-type strain KIM1001) fully complemented its virulence defect, restoring virulence to wild-type levels as expected. This result also confirmed that the long series of genetic manipulations required for sequentially deleting and then restoring the seven effector genes did not cause unexpected or off-target modifications that alter virulence phenotypes (Figure 1 and Table 1).

**Figure 1.**
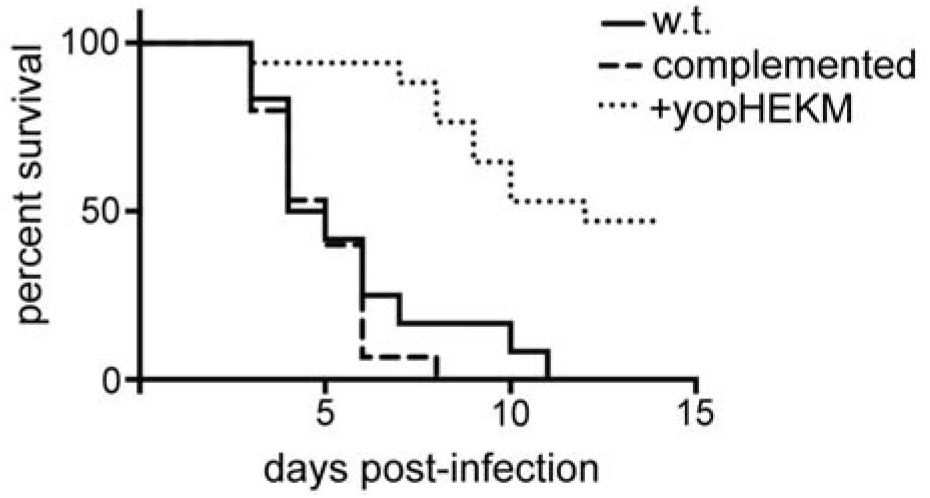
YopH, YopE, YopK, and YopM are not sufficient for full virulence. Subcutaneous infection with 1000 CFU KIM1001 (w.t.) (n=12), KIM1001ΔT3SE::+yopHEKMAJT (complemented) (n=15), or KIM1001ΔT3SE::+yopHEKM (+yopHEKM) (n=17). Mice infected with KIM1001 or with the fully complemented strain rapidly succumbed to infection, while KIM1001ΔT3SE::+yopHEKM killed 9 out of 17 mice with delayed kinetics.

### Strains expressing YopK or YopM in addition to YopH and YopE induce more effective adaptive immunity

The avirulence of strains KIM1001ΔT3SE::+yopHE, KIM1001ΔT3SE::+yopHE**M**, and KIM1001ΔT3SE::+yopHE**K** led us to investigate their potential as live attenuated vaccines. To assess the degree of protection conferred by infection with these strains, surviving mice were challenged 28 days after initial infection with 10^3^ CFU KIM1001 s.c. Mice exposed to KIM1001ΔT3SE::+yopHE**M** or KIM1001ΔT3SE::+yopHE**K** uniformly survived the challenge, while 3 out of 10 mice exposed to KIM1001ΔT3SE::+yopHE succumbed within 14 days (Figure 2). The full protection from challenge conferred by KIM1001ΔT3SE::+yopHEK and KIM1001ΔT3SE::+yopHEM suggests that each of these strains, while nonlethal, causes infection that is sufficiently persistent to trigger robust involvement of the adaptive immune system. The partial protection provided by exposure to KIM1001ΔT3SE::+yopHE indicates a weaker or less consistent adaptive immune response, suggesting that this strain is more susceptible to clearance by the innate immune system.

**Figure 2.**
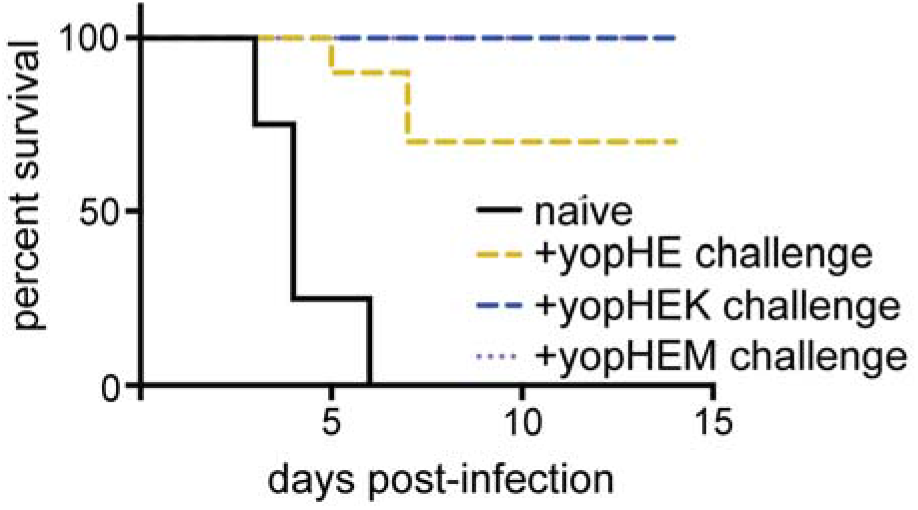
Protective immunity from exposure to *Y. pestis* mutants expressing subsets of T3SS effectors. When challenged with 1000 CFU KIM1001 28 days after initial infection, mice exposed to KIM1001ΔT3SE::+yopHEM and KIM1001ΔT3SE::+yopHEK were fully protected (n=6 mice each), while exposure to KIM1001ΔT3SE::+yopHE provided partial protection (3 of 10 mice died). Naïve mice succumbed to infection within seven days (n=8).

### YopJ, YopT, and YpkA each contribute to virulence of *Y. pestis* following subcutaneous infection

The attenuation of KIM1001ΔT3SE::+yopHEKM relative to KIM1001ΔT3SE::+yopHEKMAJT is strong evidence that at least one of the remaining effectors (YopT, YpkA, or YopJ) functionally contributes to virulence *in vivo*. However, strains deficient in any one of these effectors are not significantly attenuated (36, 37) (and see Table 1).

We generated derivatives of the KIM1001ΔT3SE::+yopHEKM strain that included a functional copy of either the *ypkA, yopT*, or *yopJ* gene in its original locus. The resulting strains were KIM1001ΔT3SE::+yopHEKM**A**, expressing YpkA; KIM1001ΔT3SE::+yopHEKM**T**, expressing YopT; and KIM1001ΔT3SE::+yopHEKM**J**, expressing YopJ. Each of these strains was substantially more virulent than KIM1001ΔT3SE::+yopHEKM. Although none caused 100% mortality, each was virulent enough that the difference between survival curves for these strains and the wild-type strain KIM1001 was not statistically significant (Figure 3 and Table 1). Given the distinct biochemical activities of YopT, YpkA, and YopJ, it is curious that each of these effectors increased virulence of the KIM1001ΔT3SE::+yopHEKM approximately equally. The different targets and activities reported for these effectors suggest that they are not truly redundant at the molecular level, and may therefore enhance virulence through distinct processes.

**Figure 3.**
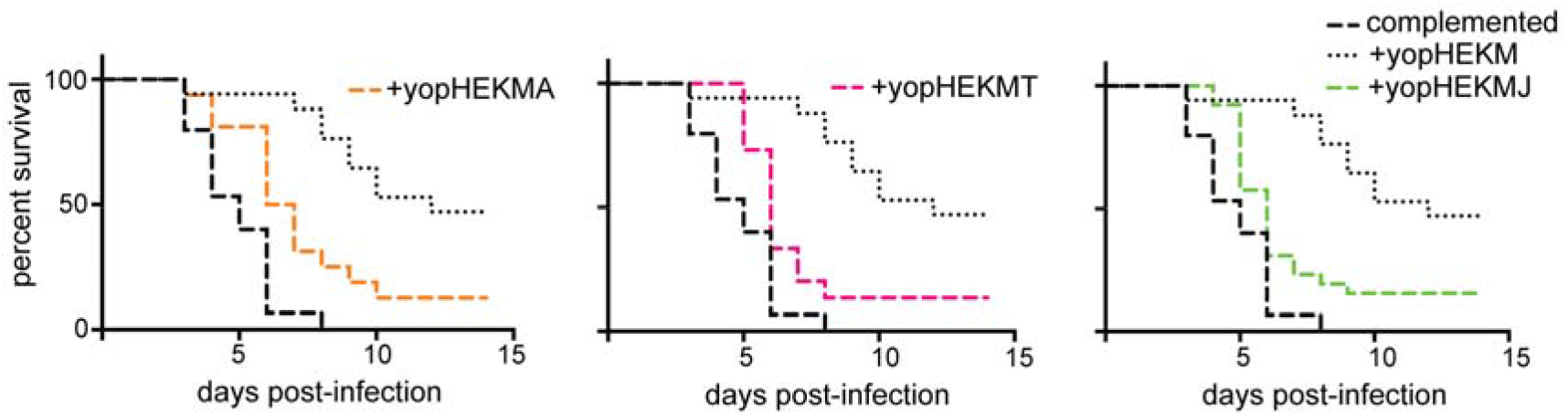
Addition of YpkA, YopT, or YopJ enhances the virulence of a *Y. pestis* strain expressing YopH, YopE, YopK, and YopM. Survival following subcutaneous infection with 1000 CFU KIM1001ΔT3 SE::+yopHEKMA (+yopHEKMA) (n=16), KIM1001ΔT3SE::+yopHEKMT (+yopHEKMT) (n=15), or KIM1001ΔT3SE::+yopHEKMJ (+yopHEKMJ) (n=26), compared to the survival curves for KIM1001ΔT3SE::+yopHEKMAJT (complemented) and KIM1001ΔT3SE::+yopHEKM (+yopHEKM) from Figure 1. KIM1001ΔT3SE::+yopHEKMA killed 14 out of 16 mice, KIM1001ΔT3SE::+yopHEKMT killed 13 out of 15 mice, and KIM1001ΔT3SE::+yopHEKMJ killed 22 out of 26 mice, all with kinetics similar to KIM1001ΔT3 SE::+y opHEKM AJT.

### YopJ suppresses immune cell recruitment in the liver

Evaluation of liver pathology following intravenous infection is a useful method to assay immune cell responses to *Y. pestis* (21, 39, 40). KIM1001ΔT3SE::+yopHEKM and the strains that additionally expressed YpkA, YopT, or YopJ were injected intravenously into mice. Livers were collected 48 hours after infection for histopathological analysis. KIM1001ΔT3SE::+yopHEKM elicited robust recruitment of immune cells, whereas KIM1001ΔT3SE::+yopHEKM**AJT**, like KIM1001, effectively suppressed accumulation of inflammatory cells at foci of bacterial growth (Figure 4A-B). The addition of YopJ to KIM1001ΔT3SE::+yopHEKM appears sufficient to fully suppress immune cell recruitment in this context, as lesions caused by KIM1001ΔT3SE::+yopHEKM**J** were indistinguishable from those caused by KIM1001. By contrast, neither KIM1001ΔT3SE::+yopHEKM**A** nor KIM1001ΔT3SE::+yopHEKM**T** suppressed inflammatory cell recruitment relative to KIM1001ΔT3SE::+yopHEKM (Figure 4A-B). YopJ, therefore, appears to be uniquely essential (though likely not sufficient) for suppressing accumulation of immune cells at sites of bacterial replication *in vivo*.

**Figure 4.**
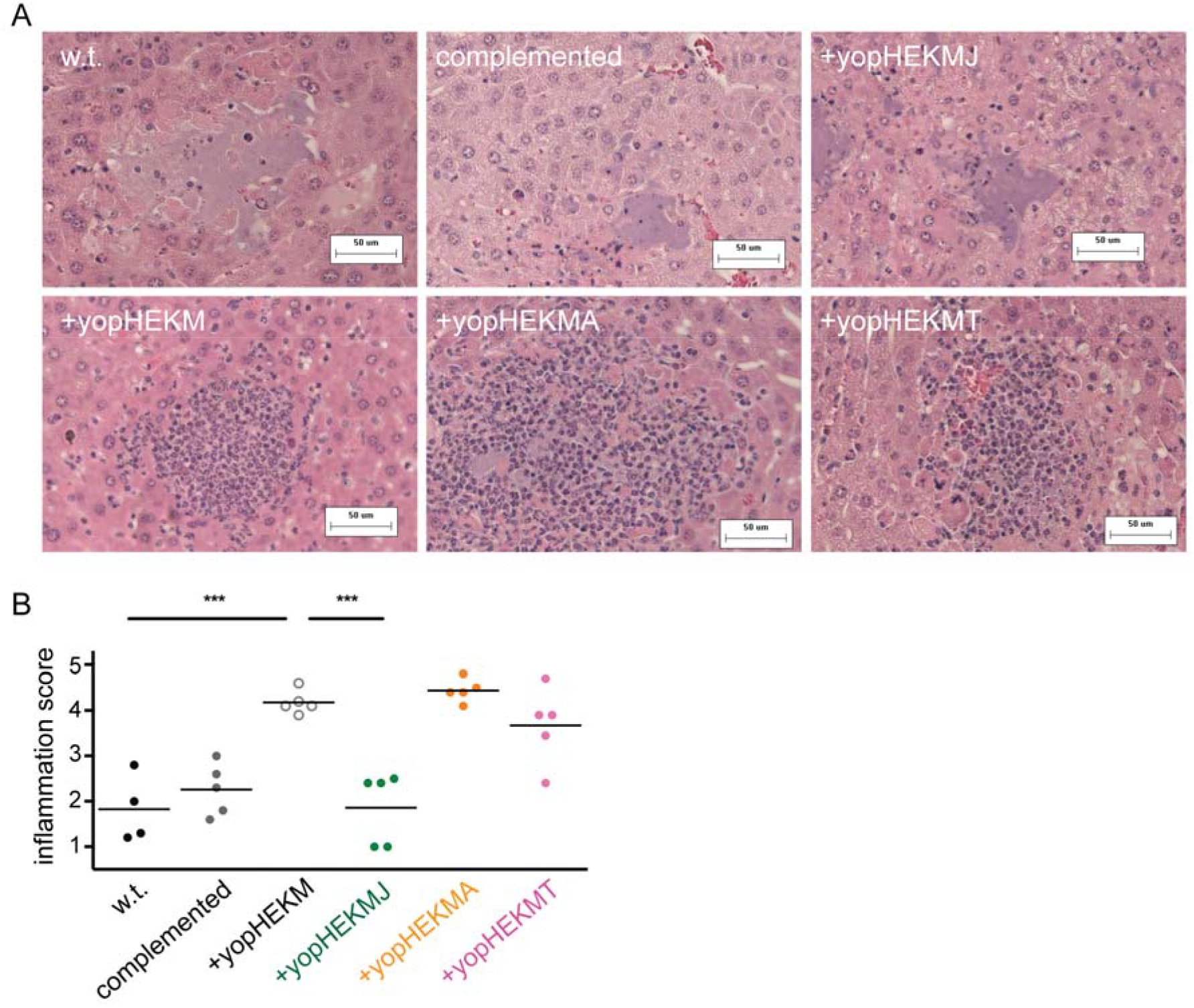
YopJ suppresses immune cell recruitment to foci of bacterial growth in liver tissue. **(A)** Representative liver sections stained with hematoxylin and eosin from mice 48 hours after intravenous infection with 10^3^ CFU KIM1001 (w.t.), KIM1001ΔT3SE::+yopHEKMAJT (complemented), KIM1001 ΔT3SE::+yopHEKMJ (+yopHEKMJ), KIM1001ΔT3SE::+yopHEKM (+yopHEKM), KIM1001ΔT3SE::+yopHEKMA (+yopHEKMA), or KIM1001ΔT3SE::+yopHEKMT (+yopHEKMT). Strains with a functional *yopJ* allele grow freely in liver tissue without attracting inflammatory cells (top row), in contrast to strains deficient in *yopJ* (bottom row). **(B)** Severity of inflammation was scored on an arbitrary scale (1 = free bacteria with few or no inflammatory cells; 5 = abundant inflammatory cells with little or no visible free bacteria). Each data point represents the average score for 10 lesions from a single mouse. Representative of two independent blinded scorings.

### YpkA and YopT are dispensable for inducing macrophage cell death and for bacterial survival in co-culture with neutrophils

The ability of the *Y. pestis* T3SS to cause apoptosis in macrophages is well established, and is considered an important function in promoting virulence. We therefore examined how various combinations of effectors influenced cell death of immortalized macrophages *in vitro*.

Some incomplete sets of *Yersinia* effectors result increased macrophage death, either via YopE induction of pyroptosis (30, 31) or as a result of dysregulated effector and translocon secretion in the absence of YopK (32, 41). However, the ΔT3SE::+yopHEKM strain expresses both YopM, which prevents YopE-mediated activation of the pyrin inflammasome (30, 31), and YopK, which prevents inflammasome activation by translocon components (41, 42). As expected, therefore, this strain resulted in minimal macrophage death (Figure S1).

Consistent with previous reports (21, 43, 44), we observed T3SS-dependent cell death induced by YopJ. The ΔT3SE::+yopHEKMJ strain caused macrophage death at a level indistinguishable from the wild-type strain (Figure S1). This YopJ-mediated cell death may contribute to the reduced visible recruitment of innate immune cells *in vivo* (Figure 4). Addition of YopT or YpkA to a strain expressing YopE, YopH, YopK, and YopM had no effect on macrophage cell death.

In addition to undermining macrophage function, *Y. pestis* must overcome the antimicrobial host responses mediated by neutrophils. The T3SS is known to be critical for evasion of neutrophil killing *in vitro* (45) (and, to some degree, *in vivo* (20)). To determine whether any of the observed synthetic phenotypes *in vivo* result from differential ability to survive neutrophil antimicrobial responses, effector mutants were assayed systematically for the ability to survive in co-culture with primary human neutrophils. To facilitate the effort of assaying a large number of bacterial strains simultaneously, we developed a luminescence-based assay to monitor bacterial survival in co-culture with human neutrophils that is higher-throughput than plating to measure colony-forming units (CFU), and has the additional benefit of providing a readout of bacterial metabolic activity in real time (see Methods). Survival of *Y. pestis* in co-culture with neutrophils required the T3SS, as expected (Figure S2). The T3SS injectisome in combination with either YopH or YopE is sufficient for this survival phenotype (Figure S3A), and deletion of both YopH and YopE together recapitulates the susceptibility to neutrophils observed in a T3SS-deficient strain (Figure S3B). No other effector is necessary (Figure S4) or sufficient (Figure S5) for bacterial survival in the presence of primary neutrophils *in vitro*. YopT and YpkA, therefore, are unlikely to enhance virulence *in vivo* by directly undermining neutrophil bactericidal activity.

## Discussion

Since the original discovery of the type III secretion system in *Y. pestis* and *Y. pseudotuberculosis* (1, 46, 47), type III secretion systems have been found to contribute to virulence not only in the *Yersiniae* but also in diverse Gram-negative pathogens, including pathogenic species of *Salmonella, Pseudomonas, Vibrio, Burkholderia*, and in enteropathogenic *Escherichia coli* strains (48–54). Multifactorial virulence determinants such as these T3SSs are crucial factors in allowing fulminant pathogens to undermine host defense systems. However, complex bacterial systems are difficult to study using traditional single-knockout methods of analysis, which are confounded by apparent functional redundancy.

In this work, we used a gain-of-function approach to determine that the four effectors previously known to be required during infection – YopH, YopE, YopK, and YopM – are not sufficient to mediate full virulence of *Y. pestis*. Indeed, strains expressing only YopH, YopE, and either YopK or YopM failed to sicken or kill any mice, although infection with these strains is protective against subsequent challenge with virulent *Y. pestis*. As these strains retain the structural components of the T3SS, as well as all other non-T3SS virulence factors, they may prove to useful in the construction of live attenuated vaccines.

YopJ, YopT, and YpkA had not been shown to contribute uniquely to *Y. pestis* virulence in mammalian infection (36, 37) (and see Table 1). We initially suspected that the apparent dispensability of these effectors was due to functional redundancy between two of them, but we found instead that all three individually increase virulence when introduced to the attenuated strain expressing only YopH, YopE, YopK, and YopM. It is possible that the action of each of these three effectors contributes additively to virulence of *Y. pestis*, and that addition of any of them to the core effector set of YopH, YopE, YopK, and YopM is sufficient to pass some threshold of immune subversion that allows robust dissemination and proliferation of *Y. pestis in vivo*.

YopJ, YopT, and YpkA increase lethality approximately equally, but have different biochemical activities and targets from one another. Strains expressing each of these effectors also behave differently in more targeted assays, supporting the model that these effectors contribute to pathogenesis independently rather than redundantly. For example, addition of YopJ to the KIM1001ΔT3SE::+yopHEKM construct blocks accumulation of inflammatory cells at foci of infection in deep tissue, while addition of either YopT or YpkA to KIM1001ΔT3SE::+yopHEKM does not produce any obvious histological signature (Figure 4). This is consistent with a recent finding by our colleagues Ratner *et al*., who report a similar phenotype in a YopJ single deletion strain (21). Ratner *et al*. also demonstrate that, when YopM is absent, YopJ is necessary for full virulence of *Y. pestis* following subcutaneous infection. The YopM-independent role for YopJ in virulence that we report here is a novel finding.

Although we have now established a functional role for YopT and YpkA in infection, we cannot yet explain how these effectors enhance the virulence of the KIM1001ΔT3SE::+yopHEKM strain. T3SS induction of macrophage cell death, while important for *Y. pestis* pathogenesis, does not appear to require either of these effectors. The work presented here is in agreement with reports that, though NLRP3/NLRC4-mediated cell death has been shown to occur in response to the needle and translocon proteins of the JG150ΔT3SE strain (21, 44), macrophage cell death mediated by the wild-type T3SS seems to depend primarily on the activity of YopJ (34, 43). The T3SS of *Y. pestis* also targets neutrophils *in vivo* (7, 20, 55), and neutrophils are key players in controlling *Y. pestis* infection (20, 56, 57). However, our work in *Y. pestis* attributes anti-neutrophil activity primarily to YopH and YopE (Figures S3-S5), consistent with previous work focusing on *Y. pseudotuberculosis* and *Y. enterocolitica* (22, 23, 58–60).

Interestingly, both YopT and YpkA interfere with Rho signaling in mammalian cells. YopT is a cysteine protease that cleaves the prenylated moiety from small GTPases of the Rho family, including RhoA and Rac1, to reduce their activity by releasing them from the cytoplasmic membrane (61). In *Y. pseudotuberculosis*, YopT also inhibits RhoG via this mechanism. This activity synergizes with YopE inhibition of RhoG to decrease phagocytic uptake of *Yersinia* (62). YopT of *Y. enterocolitica* upregulates transcription of the anti-inflammatory GILZ protein in HeLa cells and in a monocyte cell line (63), though whether this is conserved in *Y. pestis* and functional during infection is unknown. Like YopT, YpkA inhibits Rho GTPases including RhoA, Rac1, and Rac2 (summarized in (8)). Rho GTPases are also key host targets of YopE activity. The importance of multiple effectors for deranging Rho signaling is not clear, though it is possible that differential tissue tropism or effector kinetics may play a role. Future work with the set of strains we report here may provide clues regarding the function of YopT and YpkA during infection. Promising lines of inquiry include measuring the effect of these effectors on cytokine production *in vivo*, on the kinetics of distal organ colonization following subcutaneous infection, and on the ability of *Y. pestis* to establish and maintain sufficient bacteremia to reliably infect fleas feeding on infected mammals.

In addition to refining the model of *Y. pestis* pathogenesis and creating genetic tools that will streamline further study of this T3SS in the yersiniae, this work represents a general strategy for effective and efficient genetic analysis of complex bacterial systems by unmasking the functional contributions of individual components. Genetic analysis of partially redundant systems, even those of only moderate complexity such as the seven effectors of the *Yersinia* T3SS, is difficult to perform in a comprehensive manner. Combinatorial knockouts are the traditional approach to identifying functional redundancy, but an unbiased combinatorial approach rapidly becomes unfeasible as the size of the system increases. For example, even constructing and assaying all 21 possible double knockouts of *Y. pestis* T3SS effectors is not an attractive approach, and there is no guarantee that functional redundancy is limited to only two effectors. The “bottom-up” approach we describe here, to identify effector(s) that are sufficient rather than necessary for various phenotypes, is perhaps generalizable to other complex systems. We have shown that this approach can be particularly effective when combined with candidate-based hypothesis testing, as this limits the number of combinations that must be examined. Once such strains are generated, they can be assayed for multiple phenotypes in both *in vitro* and *in vivo* systems. These strain banks, therefore, allow for rapid systematic identification and disentanglement of apparent functional redundancy among components of complex bacterial systems.

## Materials and methods

### Bacterial strains and growth conditions

The genotype and source for *Y. pestis* strains are presented in Table S1. Genotype information includes the codons removed by each in-frame effector deletion: for example, the notation *yopH*^Δ3–467^ denotes that the *yopH* gene harbors an in-frame deletion of codons 3-467 (inclusive). *In vivo* experiments (lethality, liver histology) were performed using strains made in the fully virulent KIM1001 background. *In vitro* experiments (macrophage cell death and bacterial survival in the presence of neutrophils) were performed in the JG150 background (equivalent to KIM5), which lacks the *pgm* locus required for iron acquisition, to permit experimentation under biosafety level 2 conditions.

*Y. pestis* was cultured in the rich medium TB, prepared to maximize plating efficiency as previously described (64), or in the defined Serum Nutritional Medium (SNM) (18). All *Y. pestis* cultures were supplemented with 2.5 mM CaCl_2_ to suppress type III secretion. *E. coli* strains used in construction of *Y. pestis* mutants were cultured in Luria broth. Media were supplemented with 100 μg/ml ampicillin and/or 25 μg/ml diaminopimelic acid as appropriate. *Y. pestis* strains are available on request to researchers with qualifying regulatory approvals (contact).

### Construction and complementation of nonpolar mutant strains of *Y. pestis*

*Y. pestis* mutants were constructed via allelic exchange with the suicide vector pRE107 ((65), gift from D. Schifferli). Primers are listed in Table S2. Deletion mutants for each gene were constructed by amplifying flanking homology to the gene using primer pairs A+B and C+D, hybridizing the resulting fragments, and cloning this “stitched” product into the pRE107 plasmid before proceeding with allelic exchange as described (65). The *E. coli* donor strain α2155 ((66), gift from B. Akerley) was used to propagate pRE107 derivatives and to introduce them into *Y. pestis* by conjugation. Complementation of effector mutants was performed *in situ*, using pRE107-based allelic exchange to replace each deleted gene with a wild-type copy amplified using the primer pair A+D for each gene. Attenuated mutants for use in *in vitro* assays were generated by screening for spontaneous loss of the *pgm* locus on HIB agar with Congo Red, verified by PCR as described (21). Luminescent derivatives were constructed by electroporation of the pML001 plasmid containing the *lux* operon from *Photorhabdus luminescens* (67) followed by selection of transformants on media supplemented with ampicillin.

### Animal infections

All animal infections were conducted in conformity with the Guide for the Care and Use of Laboratory Animals of the National Institutes of Health, and with the review and approval of the UMass Medical School Institutional Animal Care and Use Committee (IACUC). C57BL/6 mice were infected as indicated. All bacterial cultures used for inoculation were grown at 37°C on TB agar (TB medium with 1.5% agar) containing 2.5 mM CaCl_2_ for one overnight prior to infection. Bacterial cells were diluted in infection-grade phosphate-buffered saline (PBS) to the desired concentration. In every case, the number of viable bacteria present in each dose was verified by dilution plating of the inoculum. Mice were monitored every twelve hours for signs of illness such as ruffled fur, shallow breathing, limping, reluctance to move, and swollen lymph nodes.

### Histological analysis of T3SS mutants in liver tissue

Mice were sacrificed 48 hours following intravenous infection with 10^3^ CFUs. Livers were removed, fixed in 10% neutral-buffered formalin, and embedded in paraffin for sectioning and staining. Samples were randomized and 10 lesions from each mouse were scored, blinded, for severity of inflammation in bacterial lesions. The following scale was used for scoring: 1 = free bacteria with few or no inflammatory cells; 2 = some inflammatory cells present, but free bacteria fill the majority of the lesion area; 3 = lesion area is split approximately equally between inflammatory cells and bacteria; 4 = some free bacteria are visible, but inflammatory cells fill the majority of the lesion area; 5 = abundant inflammatory cells with few or no visible free bacteria. Scoring was performed twice on scrambled, blinded samples to ensure results were consistent.

### Macrophage cell death experiments

Macrophages were monitored for cell death by ethidium homodimer (EthD1) fluorescence, as described (21). Briefly, 8x10^4^ immortalized murine macrophages from C57BL/6 mice (a gift from K. Fitzgerald) were infected with 8x10^4^ CFU of various *Y. pestis* strains grown to mid-log phase in SNM supplemented with 2.5 mM CaCl_2_. Macrophages and bacteria were added to flat-bottomed 96-well plates with black sides in DMEM supplemented with 10% ΔFBS, 10 mM HEPES, and 2 μM ethidium homodimer. Plates were centrifuged at 400 rpm for five minutes, sealed, and incubated at 37°C in a Synergy H4 microplate reader to monitor ethidium homodimer fluorescence (645 nm emission, 530 nm excitation). Ethidium homodimer uptake data was analyzed by calculating the area under the curve (AUC) for the increasing fluorescence signal and subtracting the AUC of control wells containing uninfected macrophages.

### Survival of T3SS in co-culture with primary human neutrophils

Viability assays were performed with bioluminescent *Y. pestis* strains using the plasmid pML001, which encodes the *lux* operon from *Photorhabdus luminescens* (67). Luminescence from this system requires the reduced flavin mononucleotide FMNH_2_ as a cofactor (68). As reduced FMNH_2_ is rapidly depleted if metabolism or the cell membrane is disrupted, bioluminescence in this system serves as a proxy for determining viability of the bacterial population in real time. CFU plating confirmed that the decreased luminescence of a T3SS-deficient strain after 4 hours in the presence of neutrophils (Figure S2) corresponded to a 50-80% reduction in bacterial viability compared to the media-only (no neutrophil) condition.

Whole blood was collected from healthy adult human volunteers in compliance with protocols reviewed and approved by the University of Massachusetts Medical School Institutional Review Board (IRB). Neutrophils were isolated from whole blood on a gelatin gradient as described (69). *Y. pestis* viability assays were performed in opaque white flat-bottomed 96-well plates that were coated for 1 hour with 10 μg/mL fibrinogen in phosphate-buffered saline (PBS) and washed twice with PBS. 5x10^5^ neutrophils were infected with luminescent strains of *Y. pestis* at MOI 0.1 in SNM supplemented with 2.5 mM CaCl_2_, 100 μg/ml ampicillin, and 4% normal human serum. Plates were centrifuged at 400 rpm for five minutes, sealed, and incubated at 37°C in a Synergy H4 microplate reader. Bacterial luminescence was monitored in for six hours. Three independent bacterial cultures were assayed for each *Y. pestis* strain in each experiment, and experiments were performed at least twice with neutrophils from different donors.

## Acknowledgements

We thank Dr. Brian Akerley, Dr. Dieter Schifferli, and Dr. Kate Fitzgerald for providing research materials; Dr. Beth McCormick, Andrew Zukauskas, and Christopher Louissaint for assistance with blood collection; Dr. Christopher Sassetti for his help in refining this manuscript; and Dr. Egil Lien for substantive and insightful discussions regarding the role of T3SS effectors in potentiating host cell death.

## References

1. Rosqvist R, Magnusson KE, Wolf-Watz H. 1994. Target cell contact triggers expression and polarized transfer of Yersinia YopE cytotoxin into mammalian cells. The EMBO Journal 13:964–972.

2. Persson C, Nordfelth R, Holmström A, Håkansson S, Rosqvist R, Wolf-Watz H. 1995. Cell-surface-bound Yersinia translocate the protein tyrosine phosphatase YopH by a polarized mechanism into the target cell. Molecular microbiology 18:135–150.

3. Håkansson S, Schesser K, Persson C, Galyov EE, Rosqvist R, Homblé F, Wolf-Watz H. 1996. The YopB protein of Yersinia pseudotuberculosis is essential for the translocation of Yop effector proteins across the target cell plasma membrane and displays a contact-dependent membrane disrupting activity. The EMBO journal 15:5812–5823.

4. Plano GV, Schesser K. 2013. The Yersinia pestis type III secretion system: Expression, assembly and role in the evasion of host defenses. Immunologic Research 57:237–245.

5. Edgren T, Forsberg Å, Rosqvist R, Wolf-Watz H. 2012. Type III secretion in Yersinia: Injectisome or not? PLoS Pathogens 8:2–4.

6. Akopyan K, Edgren T, Wang-Edgren H, Rosqvist R, Fahlgren A, Wolf-Watz H, Fallman M. 2011. Translocation of surface-localized effectors in type III secretion. Proceedings of the National Academy of Sciences 108:1639–1644.

7. Marketon MM, DePaolo RW, DeBord KL, Jabri B, Schneewind O. 2005. Plague bacteria target immune cells during infection. Science 309:1739–1741.

8. Pha K, Navarro L. 2016. Yersinia type III effectors perturb host innate immune responses. World Journal of Biological Chemistry 7:1–13.

9. Portnoy DA, Blank HF, Kingsbury DT, Falkow S. 1983. Genetic analysis of essential plasmid determinants of pathogenicity in Yersinia pestis. The Journal of Infectious Diseases 148:297–304.

10. Goguen JD, Yother J, Straley SC. 1984. Genetic analysis of the low calcium response in Yersinia pestis mu d1 (Ap lac) insertion mutants. Journal of Bacteriology 160:842–848.

11. Cornelis GR, Wolf-Watz H. 1997. The Yersinia Yop virulon: a bacterial system for subverting eukaryotic cells. Molecular Microbiology 23:861–867.

12. Allaoui A, Schulte R, Cornelis GR. 1995. Mutational analysis of the Yersinia enterocolitica virC operon: characterization of yscE, F, G, I, J, K required for Yop secretion and yscH encoding YopR. Molecular Microbiology 18:343–355.

13. Bozue J, Cote CK, Webster W, Bassett A, Tobery S, Little S, Swietnicki W. 2012. A Yersinia pestis YscN ATPase mutant functions as a live attenuated vaccine against bubonic plague in mice. FEMS Microbiology Letters 332:113–121.

14. Straley SC, Cibull ML. 1989. Differential clearance and host-pathogen interactions of YopE- and YopK- YopL- Yersinia pestis in BALB/c mice. Infection and Immunity 57:1200–1210.

15. Kerschen EJ, Cohen DA, Kaplan AM, Straley SC. 2004. The plague virulence protein YopM targets the innate immune response by causing a global depletion of NK cells. Infection and Immunity 72:4589–4602.

16. Bubeck SS, Dube PH. 2007. Yersinia pestis CO92 Δ yopH is a potent live, attenuated plague vaccine. Clinical and Vaccine Immunology 14:1235–1238.

17. Cantwell AM, Bubeck SS, Dube PH. 2010. YopH inhibits early pro-inflammatory cytokine responses during plague pneumonia. BMC Immunology 11:29–29.

18. Palace SG, Proulx MK, Lu S, Baker RE, Goguen JD. 2014. Genome-wide mutant fitness profiling identifies nutritional requirements for optimal growth of Yersinia pestis in deep tissue. mBio 5:e01385–01314.

19. Peters KN, Dhariwala MO, Hughes Hanks JM, Brown CR, Anderson DM. 2013. Early apoptosis of macrophages modulated by injection of Yersinia pestis YopK promotes progression of primary pneumonic plague. PLoS Pathogens 9:e1003324.

20. Ye Z, Kerschen EJ, Cohen DA, Kaplan AM, Van Rooijen N, Straley SC. 2009. Gr1+ cells control growth of YopM-negative Yersinia pestis during systemic plague. Infection and Immunity 77:3791–3806.

21. Ratner D, Orning MPA, Starheim KK, Marty-Roix R, Proulx MK, Goguen JD, Lien E. 2016. Manipulation of IL-1 β and IL-18 production by Yersinia pestis effectors YopJ and YopM and redundant impact on virulence. Journal of Biological Chemistry 291:jbc.M115.697698–jbc.M697115.697698.

22. Songsungthong W, Higgins MC, Rolán HG, Murphy JL, Mecsas J. 2010. ROS-inhibitory activity of YopE is required for full virulence of Yersinia in mice. Cellular Microbiology 12:988–1001.

23. Rolan HG, Durand EA, Mecsas J. 2013. Identifying Yersinia YopH-targeted signal transduction pathways that impair neutrophil responses during in vivo murine infection. Cell Host and Microbe 14:306–317.

24. Rosqvist R, Bolin I, Wolf-Watz H. 1988. Inhibition of phagocytosis in Yersinia pseudotuberculosis: A virulence plasmid-encoded ability involving the Yop2b protein. Infection and Immunity 56:2139–2143.

25. Rosqvist R, Forsberg A, Rimpilainen M, Bergman T, Wolf-Watz H. 1990. The cytotoxic protein YopE of Yersinia obstructs the primary host defence. Molecular Microbiology 4:657–667.

26. Fallman M, Andersson K, Hakansson S, Magnusson KE, Stendahl O, Wolf-Watz H. 1995. Yersinia pseudotuberculosis inhibits Fc receptor-mediated phagocytosis in J774 cells. Infection and Immunity 63:3117–3124.

27. Persson C, Nordfelth R, Andersson K, Forsberg Å, Wolf-Watz H, Fällman M. 1999. Localization of the Yersinia PTPase to focal complexes is an important virulence mechanism. Molecular Microbiology 33:828–838.

28. LaRock CN, Cookson BT. 2012. The Yersinia virulence effector YopM binds caspase-1 to arrest inflammasome assembly and processing. Cell Host and Microbe 12:799–805.

29. Chung LK, Philip NH, Schmidt VA, Koller A, Strowig T, Flavell RA, Brodsky IE, Bliska JB. 2014. IQGAP1 is important for activation of caspase-1 in macrophages and is targeted by Yersinia pestis type III effector YopM. mBio 5:e01402–01414.

30. Ratner D, Orning MP, Proulx MK, Wang D, Gavrilin MA, Wewers MD, Alnemri ES, Johnson PF, Lee B, Mecsas J, Kayagaki N, Goguen JD, Lien E. 2016. The Yersinia pestis Effector YopM Inhibits Pyrin Inflammasome Activation. PLoS Pathogens 12:e1006035.

31. Chung LK, Park YH, Zheng Y, Brodsky IE, Hearing P, Kastner DL, Chae JJ, Bliska JB. 2016. The Yersinia virulence factor YopM hijacks host kinases to inhibit Type III effector-triggered activation of the pyrin inflammasome. Cell Host and Microbe 20:296–306.

32. Dewoody R, Merritt PM, Houppert AS, Marketon MM. 2011. YopK regulates the Yersinia pestis type III secretion system from within host cells. Mol Microbiol 79:1445–1461.

33. Lilo S, Zheng Y, Bliska JB. 2008. Caspase-1 activation in macrophages infected with Yersinia pestis KIM requires the type III secretion system effector YopJ. Infection and Immunity 76:3911–3923.

34. Weng D, Marty-Roix R, Ganesan S, Proulx MK, Vladimer GI, Kaiser WJ, Mocarski ES, Pouliot K, Chan F, Kelliher MA, Harris PA, Bertin J, Gough PJ, Shayakhmetov DM, Goguen JD, Fitzgerald KA, Silverman N, Lien E. 2014. Caspase-8 and RIP kinases regulate bacteria-induced innate immune responses and cell death. Proceedings of the National Academy of Sciences 111:7391–7396.

35. Philip NH, Dillon CP, Snyder AG, Fitzgerald P, Wynosky-Dolfi MA, Zwack EE, Hu B, Fitzgerald L, Mauldin EA, Copenhaver AM, Shin S, Wei L, Parker M, Zhang J, Oberst A, Green DR, Brodsky IE. 2014. Caspase-8 mediates caspase-1 processing and innate immune defense in response to bacterial blockade of NF-κ B and MAPK signaling. Proceedings of the National Academy of Sciences 111:7385–7390.

36. Zauberman A, Cohen S, Mamroud E, Flashner Y, Tidhar A, Ber R, Elhanany E, Shafferman A, Velan B. 2006. Interaction of Yersinia pestis with macrophages: limitations in YopJ-dependent apoptosis. Infection and Immunity 74:3239–3250.

37. Brodsky IE, Medzhitov R. 2008. Reduced secretion of YopJ by Yersinia limits in vivo cell death but enhances bacterial virulence. PLoS Pathogens 4:e1000067.

38. Sodeinde OA, Subrahmanyam YV, Stark K, Quan T, Bao Y, Goguen JD. 1992. A surface protease and the invasive character of plague. Science 258:1004–1007.

39. Degen JL, Bugge TH, Goguen JD. 2007. Fibrin and fibrinolysis in infection and host defense. Journal of Thrombosis and Haemostasis 5:24–31.

40. Montminy SW, Khan N, McGrath S, Walkowicz MJ, Sharp F, Conlon JE, Fukase K, Kusumoto S, Sweet C, Miyake K, Akira S, Cotter RJ, Goguen JD, Lien E. 2006. Virulence factors of Yersinia pestis are overcome by a strong lipopolysaccharide response. Nature Immunology 7:1066–1073.

41. Zwack EE, Snyder AG, Wynosky-Dolfi MA, Ruthel G, Philip NH, Marketon MM, Francis MS, Bliska JB, Brodsky IE. 2015. Inflammasome activation in response to the Yersinia type III secretion system requires hyperinjection of translocon proteins YopB and YopD. MBio 6:e02095–02014.

42. Zwack EE, Feeley EM, Burton AR, Hu B, Yamamoto M, Kanneganti TD, Bliska JB, Coers J, Brodsky IE. 2017. Guanylate Binding Proteins Regulate Inflammasome Activation in Response to Hyperinjected Yersinia Translocon Components. Infect Immun 85.

43. Monack DM, Mecsas J, Ghori N, Falkow S. 1997. Yersinia signals macrophages to undergo apoptosis and YopJ is necessary for this cell death. Proceedings of the National Academy of Sciences 94:10385–10390.

44. Brodsky IE, Palm NW, Sadanand S, Ryndak MB, Sutterwala FS, Flavell RA, Bliska JB, Medzhitov R. 2010. A Yersinia effector protein promotes virulence by preventing inflammasome recognition of the type III secretion system. Cell Host and Microbe 7:376–387.

45. Spinner JL, Cundiff JA, Kobayashi SD. 2008. Yersinia pestis type III secretion system-dependent inhibition of human polymorphonuclear leukocyte function. Infection and Immunity 76:3754–3760.

46. Michiels T, Wattiau P, Brasseur R, Ruysschaert JM, Cornelis G. 1990. Secretion of Yop proteins by yersiniae. Infection and Immunity 58:2840–2849.

47. Salmond GPC, Reeves PJ. 1993. Membrance traffic wardens and protein secretion in Gram-negative bacteria. Trends in Biochemical Sciences 18:7–12.

48. Galán JE. 1996. Molecular genetic bases of Salmonella entry into host cells. Molecular Microbiology 20:263–271.

49. Shea JE, Hensel M, Gleeson C, Holden DW. 1996. Identification of a virulence locus encoding a second type III secretion system in Salmonella typhimurium. Proceedings of the National Academy of Sciences 93:2593–2597.

50. Frithz-Lindsten E, Du Y, Rosqvist R, Forsberg A. 1997. Intracellular targeting of exoenzyme S of Pseudomonas aeruginosa via type III-dependent translocation induces phagocytosis resistance, cytotoxicity and disruption of actin microfilaments. Molecular Microbiology 25:1125–1139.

51. Park KS, Ono T, Rokuda M, Jang MH, Okada K, Iida T, Honda T. 2004. Functional characterization of two type III secretion systems of Vibrio parahaemolyticus. Infection and Immunity 72:6659–6659.

52. Tam VC, Serruto D, Dziejman M, Brieher W, Mekalanos JJ. 2007. A type III secretion system in Vibrio cholerae translocates a formin/spire hybrid-like actin nucleator to promote intestinal colonization. Cell Host and Microbe 1:95–107.

53. Winstanley C, Hales BA, Hart CA. 1999. Evidence for the presence in Burkholderia pseudomallei of a type III secretion system-associated gene cluster. Journal of Medical Microbiology 48:649–656.

54. Jarvis KG, Kaper JB. 1996. Secretion of extracellular proteins by enterohemorrhagic Escherichia coli via a putative type III secretion system. Infection and Immunity 64:4826–4829.

55. Pechous RD, Sivaraman V, Price PA, Stasulli NM, Goldman WE. 2013. Early host cell targets of Yersinia pestis during primary pneumonic plague. PLoS Pathogens 9:e1003679.

56. Lukaszewski RA, Kenny DJ, Taylor R, Rees GC, Hartley MG, Oyston PCF, Rees DGC. 2005. Pathogenesis of Yersinia pestis infection in BALB/c Mice: effects on host macrophages and neutrophils. Infection and Immunity 73:7142-7150.

57. Laws TR, Davey MS, Titball RW, Lukaszewski R. 2010. Neutrophils are important in early control of lung infection by Yersinia pestis. Microbes and Infection 12:331–335.

58. Andersson K, Magnusson KE, Majeed M, Stendahl O, Fällman M. 1999. Yersinia pseudotuberculosis-induced calcium signaling in neutrophils is blocked by the virulence effector YopH. Infection and Immunity 67:2567–2574.

59. Westermark L, Fahlgren A, Fällman M. 2014. Yersinia pseudotuberculosis efficiently escapes polymorphonuclear neutrophils during early infection, vol 82.

60. Grosdent N, Maridonneau-Parini I, Sory MP, Cornelis GR. 2002. Role of Yops and adhesins in resistance of Yersinia enterocolitica to phagocytosis. Infection and Immunity 70:4165–4176.

61. Shao F, Dixon JE. 2003. YopT Is a Cysteine Protease Cleaving Rho Family GTPases, p 79–84. *In* Skurnik M, Bengoechea JA, Granfors K (ed), The Genus Yersinia, vol 529. Springer US, Boston, MA.

62. Mohammadi S, Isberg RR. 2009. Yersinia pseudotuberculosis virulence determinants invasin, YopE, and YopT modulate RhoG activity and localization. Infection and Immunity 77:4771–4782.

63. Köberle M, Göppel D, Grandl T, Gaentzsch P, Manncke B, Berchtold S, Müller S, Lüscher B, Asselin-Labat M-L, Pallardy M, Sorg I, Langer S, Barth H, Zumbihl R, Autenrieth IB, Bohn E. 2012. Yersinia enterocolitica YopT and Clostridium difficile toxin B induce expression of GILZ in epithelial cells. PloS One 7:e40730–e40730.

64. Pouliot K, Pan N, Wang S, Lu S, Lien E, Goguen JD. 2007. Evaluation of the role of LcrV-Toll-like receptor 2-mediated immunomodulation in the virulence of Yersinia pestis. Infection and Immunity 75:3571–3580.

65. Edwards RA, Keller LH, Schifferli DM. 1998. Improved allelic exchange vectors and their use to analyze 987P fimbria gene expression. Gene 207:149–157.

66. Dehio C, Meyer M. 1997. Maintenance of broad-host-range incompatibility group P and group Q plasmids and transposition of Tn5 in Bartonella henselae following conjugal plasmid transfer from Escherichia coli. Journal of Bacteriology 179:538–540.

67. Pan NJ, Brady MJ, Leong JM, Goguen JD. 2009. Targeting type III secretion in Yersinia pestis. Antimicrobial Agents and Chemotherapy 53:385–392.

68. Meighen EA. 1991. Molecular biology of bacterial bioluminescence. Microbiological Reviews 55:123–142.

69. Henson PM, Oades ZG. 1975. Stimulation of human neutrophils by soluble and insolubleimmunoglobulin aggregates. Journal of Clinical Investigation 56:1053–1061.

